# Refining Bulk Segregant Analyses: Ontology-Mediated Discovery of Flowering Time Genes in *Brassica oleracea*

**DOI:** 10.1101/2021.08.11.455982

**Authors:** Rutger A. Vos, Catharina A. M. van der Veen-van Wijk, M. Eric Schranz, Klaas Vrieling, Peter G. L. Klinkhamer, Frederic Lens

## Abstract

**Background:** Bulk segregant analysis (BSA) can help identify quantitative trait loci (QTLs), but this may result in substantial bycatch of functionally irrelevant genes. Here we develop a Gene Ontology-mediated approach to zoom in on specific markers implicated in flowering time from among QTLs identified by BSA of the giant woody Jersey kale phenotyped in four bulks of flowering onset.

**Results:** Our BSA yielded tens of thousands of candidate genes. We reduced this by two orders of magnitude by focusing on genes annotated with terms contained within relevant subgraphs of the Gene Ontology. A further enrichment test led to the pathway for circadian rhythm in plants. The genes that enriched this pathway are attested from previous research as regulating flowering time. Some of these genes were also identified as having functionally significant variation compared to *Arabidopsis*

**Conclusions:** We validated and confirmed our ontology-mediated results through a more targeted, homology-based approach. However, our ontology-mediated approach produced additional genes of putative importance, showing that the approach aids in exploration and discovery. We view our method as potentially applicable to the study of other complex traits and therefore make our workflows available as open-source code and a reusable Docker container.

## Background

Identifying the genes that underlie quantitative trait variation is one of the main challenges in genetics and, to the extent that this is attainable *in silico*, in bioinformatics. One appealingly straightforward approach to discovering candidate loci involved in quantitative trait differences is to sort individuals of a segregating, crossed population into ‘bulks’ defined by extremes in trait values and then interrogating the genetic contrasts between these bulks, i.e. bulk segregant analysis (BSA [1, 2]). High-throughput sequencing of pooled DNA of bulks has made it possible to quickly generate haystacks of data at low cost, within which are the genetic needles (genomic regions, specific genes, and finally SNPs) that caused the salient differences between the bulks.

Several statistics have been developed to aid in the discovery of candidates of these needles. For each SNP in a sequenced bulk, metrics exist that express its relative coverage compared to other bulks (the Δ(SNP-index) *sensu* Takagi et al. [3]) or whether its allele frequency deviates from the expectation (the modified G’ statistic of Magwene et al. [4]). Then, having defined a threshold value for the metric and using a sliding window approach, regions of (more or less) contiguous SNPs in whose metric values the bulks differ can be found, resulting in putative quantitative trait loci (QTLs) in the form of genomic regions. If the analysis is performed using a sufficiently annotated reference genome to map against, genes and SNPs that regulate the trait and intersect with the intervals should be directly locatable. However, this general approach is somewhat imprecise (and more so with low thresholds or large window sizes), resulting in a lot of ‘bycatch’ of irrelevant genes. Here, we present an approach to remove such bycatch and obtain more refined result sets by traversing and pruning subgraphs of Gene Ontology [5] annotations and KEGG pathways [6] enriched by the initial QTL finding. We apply and validate this approach using bulks determined by contrasting flowering time in a *Brassica oleracea* cross.

*B. oleracea* (family Brassicaceae, tribe Brassiceae) is a diploid plant species that gave rise to many vegetable crops rich in essential nutrients such as vitamin C and anti-carcinogens such as sulforaphane [7]. The human-induced selection for diverse edible parts of *B. oleracea* has resulted in a remarkable phenotypic diversity that exceeds the variation in most wild species. Based on the specialized morphology of the edible structures, the species is often classified into kales (leaves), cabbages (leaves), cauliflower (inflorescences), broccoli (inflorescences), Brussels sprouts (axillary buds), and kohlrabi (stems and leaves). Furthermore, *B. oleracea* (n= 9), together with its close relative *B. rapa* (n = 10), are the parental species that gave rise to *B. napus* (n = 19), arguably the most economically valuable species of the tribe Brassicaceae (canola or rapeseed).

Research in *Brassica* has been advanced by the release of genomes from various species (e.g. [8–10]), including two reference genomes from *B. oleracea*: the rapid cycling line TO1000DH3 [11] and *B. oleracea* var. *capitata* [12]. All the *Brassica* genomes show evidence of a Brassicaceae lineage-specific whole-genome triplication since their divergence from the *Arabidopsis* lineage [13], followed by diploidization that involved substantial genome reshuffling and gene losses [8, 14, 15]. This makes *Brassica* an ideal model to study polyploid genome evolution, duplicated gene loss mechanisms, neo- and subfunctionalization, and its associated impact on the vast intraspecific morphological diversity [16].

A critical agronomic trait that shows remarkable variation in *B. oleracea* is flowering time. For example, whereas broccoli is a short-lived annual that flowers in the year it was planted, cabbage is biannual, needing a cold period to induce flowering (i.e. vernalization) [17]. The genetics that underlies variation in flowering time within and among *Brassica* species is reasonably well characterized [18]. The FLOWERING LOCUS T (FT) locus, its transcriptional repressor FLOWERING LOCUS C (FLC), and its transcriptional activator CONSTANS (CO) all play a central role both in *B. oleracea, B. rapa* and in *Arabidopsis*. However, the way FT expression is modulated differs between *Brassica* and *Arabidopsis*. The overall flowering time pathway is much more complex in all cases, involving over two dozen other genes in multiple, divergent copies scattered across the genome [18]. As the exact locations of these copies are mostly known, sufficient background information is available to validate and interpret the results of the analysis we present here and assess its potential for applicability in less well-characterized traits. To provide further context for our results, we also resequenced the novel genome of the late-flowering, heterozygous, giant woody walking stick kale native to Jersey Island (cultivar *B. oleracea* convar. *acephala* var. *viridis*), one of the two parents of the BSA crosses (the other being the rapid cycling, homozygous line TO1000DH3, which has been sequenced before [11]). For this Jersey kale genome, we assess the impact of SNPs within known flowering time pathway genes and compare and contrast these with the genes discovered through our BSA analysis.

## Methods

### Plant material, crosses, genotyping and phenotyping

We crossed the homozygous doubled haploid *B. oleracea* kale-like alboglabra line TO1000DH3 [11] with the giant woody walking stick kale (*B. oleracea* convar. *acephala* var. *viridis*) native to Jersey (Channel Islands, UK [19, 20]), the latter grown from seeds ordered from Mr and Mrs Johnson, who own a company making artisanal walking sticks (Homestill, La Grande Route de St. Jean, St. Helier, Jersey, Channel Islands). We selected TO1000DH3 for its rapid flowering time and short generation time (approx. 65 days). In contrast, the Jersey kale is extremely late flowering, has a much longer generation time (at least six months), and requires a vernalization period. We crossed the two parents reciprocally, resulting in F1 seeds from both parents, which we established in tissue culture and potted in soil. We genotyped this F1 population with an allele-specific assay (KASP) on our in-house high throughput SNP genotyping platform and phenotyped the plants on time till first flowering based on two individuals per genotype, distinguishing early (EF), intermediate (IF), late (LF) and non-flowering (NF, at time of DNA extraction). We set the boundaries between these different cohorts such that we obtained bulks of roughly equal numbers of individuals (around ten per bulk; Fig. 1) and increased phenotypic contrast between late and non-flowering accessions by skipping four especially late flowering individuals in the LF bulk.

**Figure 1.**
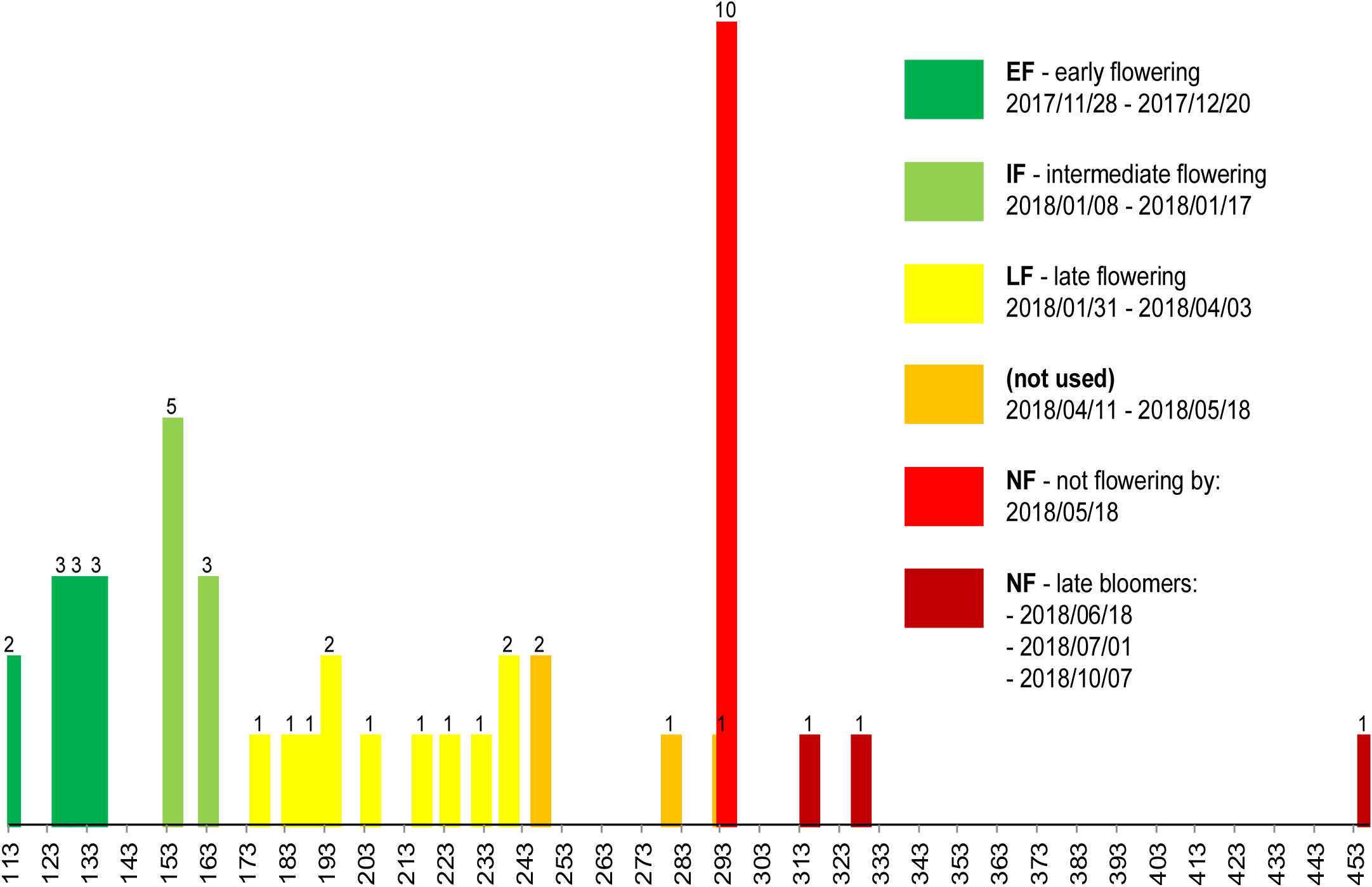
phenotyping results and assignment to bulks. F1 individuals were assigned to one of four bulks: Early Flowering (EF), Intermediate Flowering (IF), Late Flowering (LF) or Non-Flowering (NF). Of the latter, three individuals (late bloomers) flowered anyway, after DNA extraction. Four accessions were not used in order to create a greater contrast between late and non-flowering phenotypes.

### DNA extraction and sequencing data pre-processing

We performed genomic DNA extractions on a KingFisher Flex magnetic particle processor robot (Thermo Scientific) using a NucleoMag® 96 Plant kit (Macherey-Nagel GmbH & Co.). We used a volume of 150 μl for elution. We measured DNA concentrations on a Dropsense (TRINEAN NV) using a DropPlate 96-S. Based on these measurements, we pooled the DNAs of the same phenotype (EF, IF, LF and NF) equimolarly to create four DNA bulks for sequencing. We prepared libraries according to the protocol of Macrogen, containing random fragmentation of the DNA sample followed by 5’ and 3’ adapter ligation, amplification of the adapter-ligated fragments using unique index primers and gel purification. From this, we sent 400ng DNA aliquot to Macrogen for paired-end sequencing on the Illumina HiSeq X platform (read length 150bp) on a shared run. We used the BWA-MEM [21] and SAMtools [22] toolchain to map each bulk’s reads against the *B. oleracea* TO1000DH3 reference genome v2.1 of EnsemblPlants release 39, which we filtered so that we mapped against chromosomes only. We then used GATK HaplotypeCaller [23, 24] for variant (i.e. SNP and indel) calling, yielding the results summarized in Table 1.

**Table 1.** Summary results of the sequencing of bulks of early (EF), intermediate (IF), late (LF) and “non” flowering (NF, actually not flowering at time of DNA extraction) phenotypes. Bulk size refers to the number of individuals pooled for that phenotype. Coverage is given as a) total read bases divided by reference genome size; and b) average mapped coverage.

| Phenotype | Bulk size | Total read bases (bp) | Total reads | GC (%) | Q20 (%) | Q30 (%) | Coverage a, b | Variants |
| --- | --- | --- | --- | --- | --- | --- | --- | --- |
| *EF* | 11 | 60,303,831,724 | 399,363,124 | 36.93 | 95.02 | 89.25 | 123, 108 | 40,224,519 |
| *IF* | 8 | 56,804,209,216 | 376,186,816 | 36.92 | 94.84 | 88.94 | 116, 103 | 43,785,856 |
| *LF* | 11 | 54,587,863,530 | 361,509,030 | 37.14 | 96.00 | 91.34 | 112, 100 | 42,852,937 |
| *NF* | 9 | 54,296,890,456 | 359,582,056 | 37.13 | 96.84 | 92.81 | 111, 99 | 42,213,427 |

### Bulk genotyping and QTL region analysis

Given that we phenotyped the F1s by flowering time binned in four bulks, there are six pairs of contrasts (i.e. EF ↔ IF, EF ↔ LF, EF ↔ NF; IF ↔ LF, IF ↔ NF; and LF ↔ NF). We performed joint genotyping for these contrasts using the GATK CombineGVCFs/GenotypeGVCFs workflow. We then filtered these genotypes further, excluding low coverage per sample (<40x), low coverage across the pair of merged samples (<100x), unusually high coverage (>400x, e.g. repeats), low values for the GATK Genotype Quality score (<99), and low values for the frequency of the reference allele (<0.2, a conservative value as TO1000DH3 is homozygous). We calculated smoothed G statistics (G’, see [4]) over a sliding window 1Mb wide, filtering outliers by Δ(SNP-index) [3] and retaining all SNPs with G’>2.5 for further analysis. We then performed a QTL-seq analysis [3] to identify candidate QTL regions by simulation using 10k replicates and a two-sided 95% confidence interval (see Fig. 2). For the G’ and QTL-seq calculations and simulations, we used the R package QTLseqr [25]. Based on our inferred QTL regions and smoothed G’ values, we scanned the mapped assembly of each bulk for genes that fall within QTL regions and have non-synonymous SNPs with high G’. Gene coordinates were based on the annotation of the TO1000DH3 (i.e. the *B. oleracea* GFF3 release v2.1.39 of EnsemblPlants; [11]). To cross-reference the products of these genes with other information resources, we then mapped the *B. oleracea* genes to the curated and machine-predicted proteomics identifiers of UniProtKB/TrEMBL [26] using BioMart [27].

**Figure 2.**
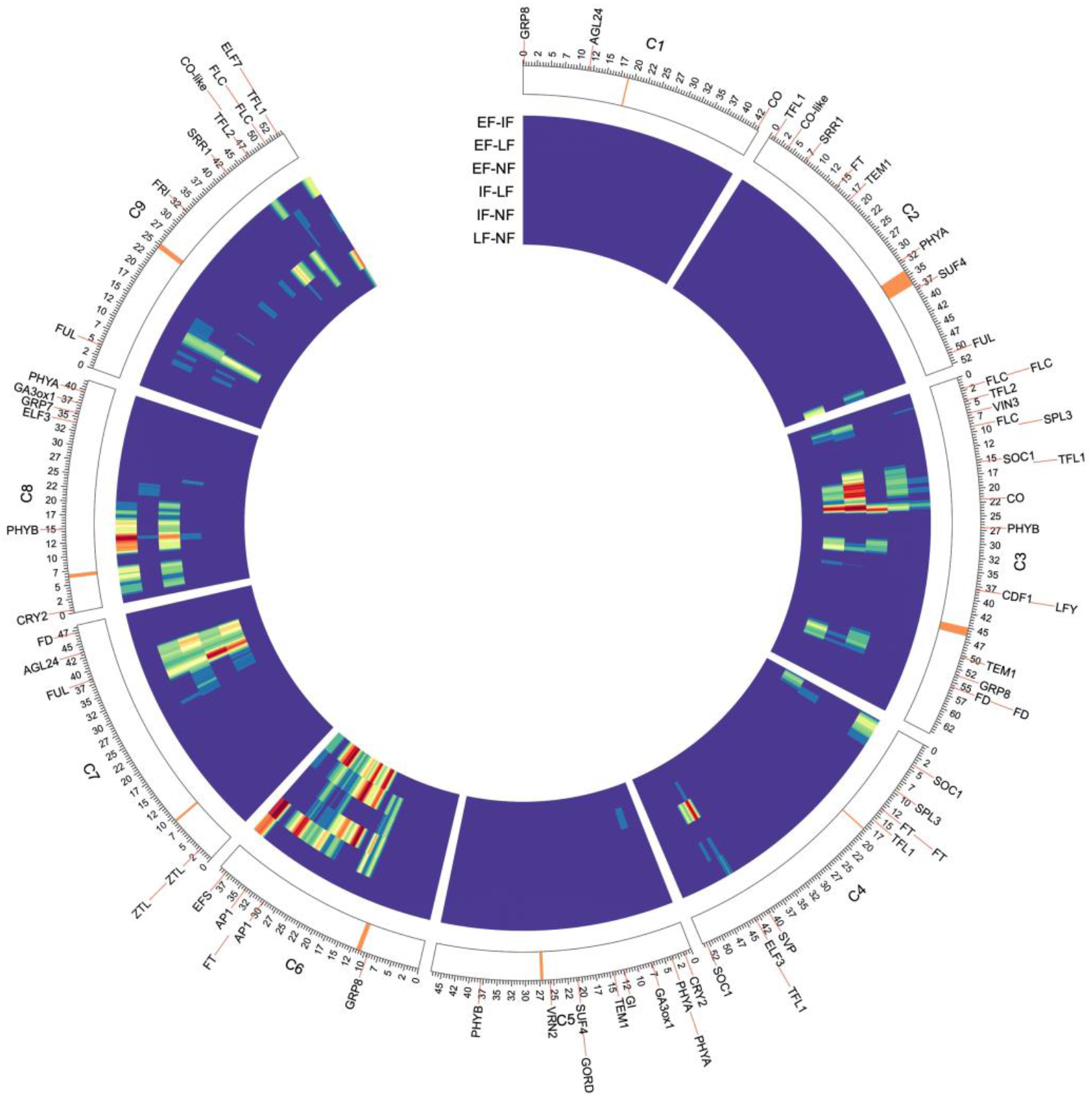
‘circos’ plot of QTL regions. Six concentric heatmaps show QTLs identified for the six possible contrasts among the four bulks. The QTLs are mapped on the annotated TO1000DH3 reference, showing the locations of flowering time genes previously identified for *B. oleracea* [18]. Centromeres indicated in orange.

### Functional enrichment, refinement, and pathway analysis

We performed singular enrichment analyses (SEA, [28]) separately for all six contrasts using the agriGO web service [29], which uses the Blast2GO [30] results for *B. oleracea* compiled by the Blast2GO Functional Annotation Repository (B2G-FAR, [31]) to establish a reference list against which to assess term enrichment by way of a hypergeometric test corrected for multiple comparisons using the Benjamini–Yekutieli method [32]. To determine the overlap between our SEAs, we merged their results in a cross-comparison (SEACOMPARE, [29]), which showed congruence in enriching numerous terms related to reproduction across all contrasts. For each of the SEA result sets, we pruned the enriched (FDR<0.05) subgraph by retaining only those terms that are reproductive developmental processes, i.e. that are subtended by the upper-level term *developmental process involved in reproduction* (GO:0003006) from the domain *biological process* of the Gene Ontology [5]. Within the pruned subgraph (Fig. 3), three out of the top-level terms are related to flower development or morphogenesis, one to seed maturation, and one (GO:0010228) is defined as:

**Figure 3.**
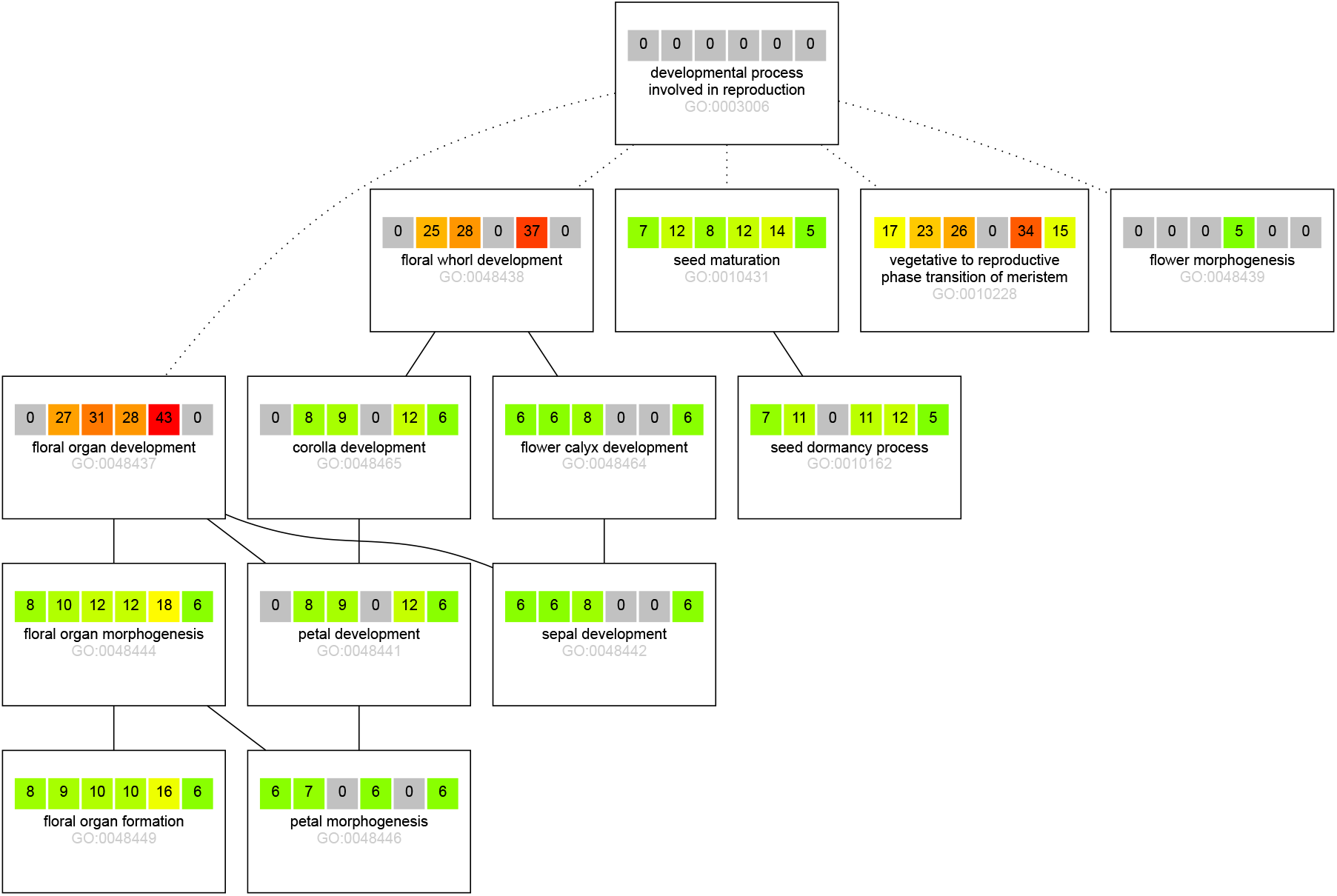
genes annotated to GO subgraph terms. Each box represents a GO term subtended by GO:0003006. Within each box, the six bulk contrasts for four bulks (i.e. EF ↔ IF, EF ↔ LF, EF ↔ NF; IF ↔ LF, IF ↔ NF; and LF ↔ NF) are shown as squares. Each square shows the number of genes annotated with that term for that contrast. The color coding corresponds with the significance of the enrichment as an inverse heatmap, i.e. the smaller *p*, the hotter the tint.

> *“The process involved in transforming a meristem that produces vegetative structures, such as leaves, into a meristem that produces reproductive structures, such as a flower or an inflorescence*.*”*

As this developmental process precedes those defined by the other top-level terms in the subgraph, we took the list of genes annotated to these terms. We used this as the input for a pathway enrichment analysis as implemented in g:Profiler [33]. This yielded an alternative view in the extent to which the genes enrich other GO terms, as well as any KEGG [6] pathways.

### Genome analysis of the Jersey kale

To gain more background insight into the genome of the giant woody Jersey kale as a potential model in general, and with an eye on differences with TO1000DH3 in flowering time loci in particular, we also sequenced the genome of a specimen of this cultivar. We followed the same protocols for DNA extraction, sequencing, genome assembly, and variant calling described for the bulks in the section *DNA extraction and sequencing data pre-processing*. However, as there was no pooling of multiple individuals (i.e. no BSA), the coverage for this single individual was commensurately higher (approx. 100x coverage). For this Jersey kale genome, we then used SnpEff [34] to assess the impact of SNPs within the loci previously identified as participating in the flowering time pathway of *Arabidopsis* [18].

## Results

The F1 seeds from our crosses resulted in 42 distinct genotypes that we successfully established in tissue culture and transferred in soil. We found a 420-day time lag between the earliest and the latest flowering F1 genotypes, presumably owing to the heterozygosity of the Jersey kale parent: the first two F1 genotypes (genotype numbers 17135, 17136) started to flower 113 days after potting, while more than a year later, at day 533, the last F1 flowered (genotype number 17109). The first bulk, early flowering EF, comprised 11 F1 genotypes that flowered between 113-135 days after potting. The second bulk, flowering at intermediate age IF, included eight F1s that flowered from 154-164 days after potting. The third bulk, late-flowering LF, represented 11 F1 genotypes that started to flower from 176-239 days after potting. The fourth bulk, non-flowering (NF) at the time of DNA extraction (day 294), included 9 F1s that only flowered after DNA extraction, up to 533 days after potting. Phenotyping results are summarized in Fig. 1 and detailed in Supplementary Table 1.

Sequencing resulted in yields per bulk between 54.3×10^9^ and 60.3×10^9^ bases, observed in 359.6×10^6^ through 399.4×10^6^ reads (which are 150bp on HiSeq X). Given the size of the reference genome (approx. 488Mbp [11]), this corresponds with a depth in the range of 111x-123x per bulk, or about 10x per individual. We retained most of the estimated raw coverage in the assemblies, yielding an average mapped coverage of 99x-108x (more sequencing statistics are listed in Supplementary Table 2). Following variant calling and joint genotyping of the six pairwise comparisons of phenotype bulks, our G’ sliding window analysis produced fairly consistent results across all comparisons. We found regions with windows of G’>2.5 on all chromosomes but C1. Still, regions that featured in the majority of pairwise comparisons were restricted to the q arm of C3 (spanning, for example, the locus of one of the CO copies), C6 (spanning one of the FT loci), C7 (spanning a FUL copy) and C8 (spanning a PHYB copy). A comparative view of these G’ regions is shown in Fig. 2. At this stage of the analysis, the QTL regions intersected with the genomic coordinates of 14,257 genes, out of which 10,469 had non-synonymous SNPs.

The term enrichment analyses (SEA, [28]) yielded between 812 and 1409 significantly enriched terms for the six bulk contrasts. Interestingly, the number of terms appears to covary somewhat with the magnitude of the trait differences, in that contrasts between the contiguous cohorts EF ↔ IF and LF ↔ NF enriched the lowest numbers of terms (1086 and 812, respectively) while those between the non-contiguous cohorts IF ↔ NF and EF ↔ NF returned the most terms (1409 and 1280). Nevertheless, the different analyses’ results overlapped extensively, as the total number of distinct terms returned overall was 1544. This was confirmed by SEACOMPARE ([29]), which also indicated extensive overlap across the six comparisons (shown in Supplementary Table 3).

Each of the comparisons enriched a subgraph of the GO topology, which we pruned further to retain only those parts subtended by GO:0003006 (*developmental process involved in reproduction*). Across the comparisons, this resulted in a consensus subgraph that spanned 14 enriched terms, shown in Fig. 3. Among these 14 terms were five upper-level terms that have an implied ordering in time and specificity with respect to their contribution to the onset and further development of flowering:

1. GO:0010228 - *vegetative to reproductive phase transition of meristem*
2. GO:0048439 - *flower morphogenesis*
3. GO:0048437 - *floral organ development*
4. GO:0048438 - *floral whorl development*
5. GO:0010431 - *seed maturation*

Terms 2-5 can only start to play a role once the transition of meristem from vegetative to reproductive has commenced (and likewise, seed maturation can only happen in a fully developed flower). Hence, we then zoomed in on only those genes that are (transitively) annotated with GO:0010228 and used these as input for g:Profiler [33], whose results are shown in Fig. 4.

**Figure 4.**
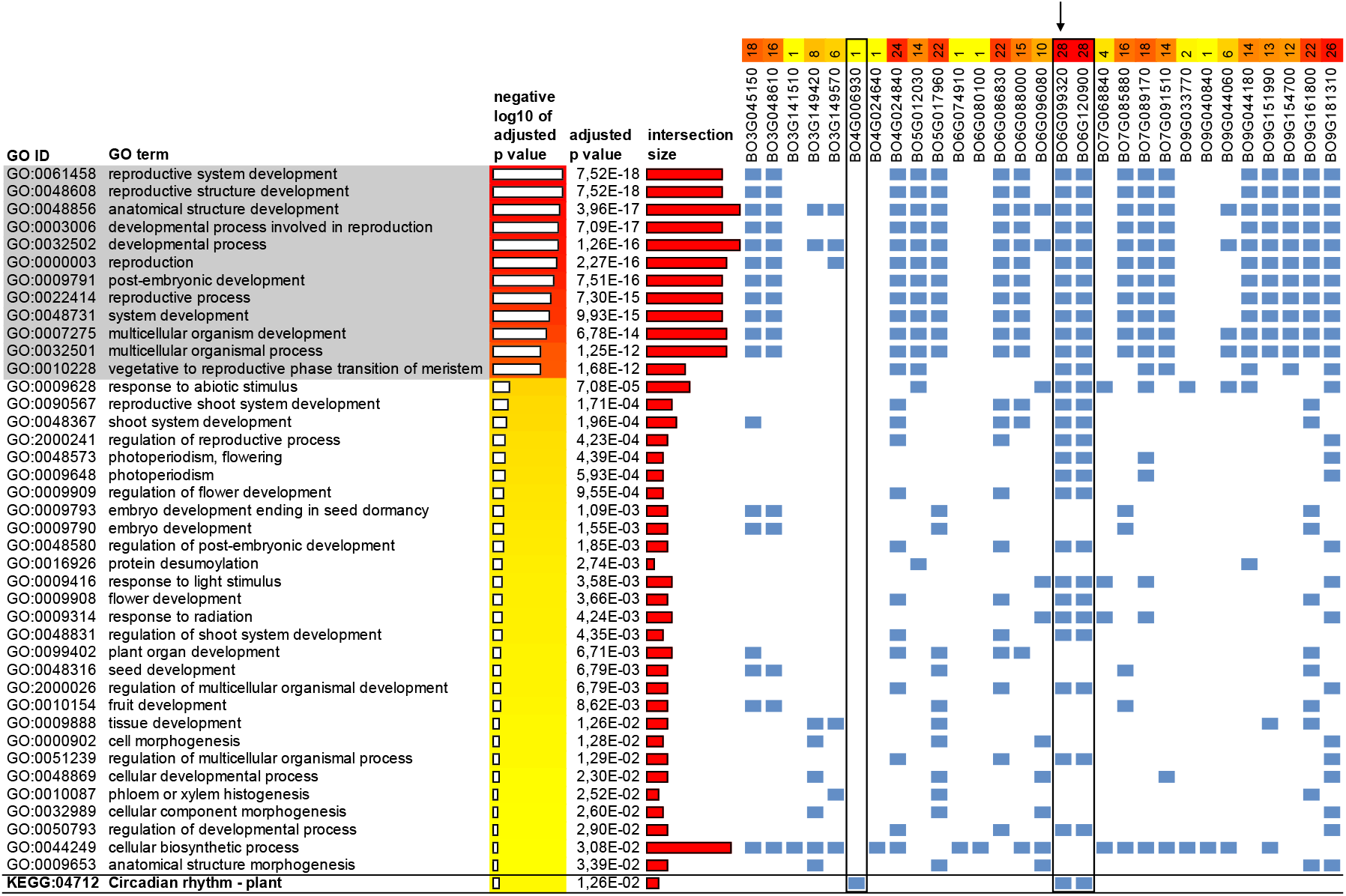
g:Profiler results for genes retained in the pruned GO subgraph. In this analysis, the genes (the columns labeled with BO gene IDs) annotated by GO:0010228 are used as input. In consequence, the GO terms above this term (both in this table and topologically in the GO graph, here shown with gray background) are heavily enriched. Below this are related terms that are also significantly enriched by the gene list. Blue cells indicate which genes contribute to the enrichment. In this analysis, the g:Profiler tool also returns enriched KEGG pathways. In this case this is only – but importantly – “KEGG:04712, Circadian rhythm – plant”.

The g:Profiler results show enrichment both for terms above GO:0010228 and those below it. The terms shown with gray background in Fig. 4 are inevitably enriched because we restricted the input gene list to those whose annotations descend from GO:0010228 – and therefore also from all upper terms ‘above’ it. More interesting are the terms below it, some of which are more specific and shed light on what, according to these annotation sets, triggers the phase transition: light and photoperiodism. However, this analysis step’s most salient result is the discovery of KEGG [6] pathway 04712 *Circadian rhythm – plant*, based on the presence of Bo4G006930 (CIRCADIAN CLOCK ASSOCIATED 1, CCA1), Bo6G099320 (FLOWERING TIME, FT) and Bo6G120900 (TWIN SISTER OF FT, TSF).

Genome sequencing of the Jersey kale yielded 1,092,319,676 forward and reverse reads (150bp). Mapped against the TO1000DH3, this resulted in an assembly covering 87.6% of the reference genome with an average depth of 170.6x. Variant calling on this assembly produced a total of about 7.5×10^6^ raw variants of all types (i.e. including indels and polymorphisms longer than 1bp). The SnpEff analysis, which assessed the impact of variants in *Arabidopsis* flowering time pathway homologs, returned results of comparable magnitude as obtained in previous research in the *B. oleracea* cultivar ‘Kashirka’ [18]. For example, most gene copies were affected by, at least, non-synonymous SNPs, and much fewer of those by splicing variation or indels causing frameshifts. Similarly to ‘Kashirka’, one copy of FT and one of FLC had indels, while the CO copies had non-synonymous SNPs but no indels. The detailed results of the analysis are available in Supplementary Table 4.

## Discussion

Our analysis was somewhat complicated by the proliferation of flowering time cohorts (four bulks, where BSA studies typically consider only the two extremes of the trait value distribution) and the commensurate increase in bulk contrasts to consider (*n*(*n*-1)/2 working out to six contrasts for four bulks). Another complication was the right censoring in waiting time till flowering for bulk NF (“non-flowering”): as three plants in this bulk flowered after the point of DNA extraction while others never did, the bulk is a mixture of very late flowering and non-flowering genotypes. Nevertheless, our results showed consistency across all comparisons in the discovered QTL regions and the GO terms the genes in these regions enriched.

An interesting result was that the extremes in the numbers of enriched GO terms loosely corresponded with the magnitude of the difference in trait values between bulks in a comparison: greater differences in flowering time enriched more terms, smaller differences fewer. For our present purposes, the increase in the number of enriched terms with greater trait differences constituted a loss in precision: as bulks are more different in flowering time, differences in the onset of contingent developmental processes (e.g. seed maturation) have a chance to manifest as well, clouding the picture and yielding more GO terms.

Our key finding was that the iterative pruning of the enriched GO subgraphs substantially reduced the number of candidate QTLs and genes in the result set. We first focused on the upper-level term GO:0003006 (*developmental process involved in reproduction*) and then zoomed in further on nodes subtended by its descendant GO:0010228 (*vegetative to reproductive phase transition of meristem*). This progression was discovered from the data and should therefore be transferrable to other systems without prior knowledge of the underlying genetics or annotations. As a result of this approach, we reduced the number of candidate genes from 10,469 to a final set of 29 genes resulting from the g:Profiler analysis. Considering that these genes include those previously established as key in regulating flowering time (both in their being homologous to those in *Arabidopsis* and in their variation between *Brassica* cultivars [18]), we view our approach as a powerful complement to existing workflows in processing BSA results.

A potential weakness is that the technique’s usefulness hinges on the quality of genome annotations and KEGG pathways: without the combination of good functional characterization (often inferred from homology) and a known background against which to perform the hypergeometric tests, gene set enrichment analyses cannot work. In practice, this means that the technique will be most applicable to well-studied model organisms or reasonably close relatives of *Arabidopsis*.

Previous research in flowering time in *B. oleracea* cultivars used the pathway in *Arabidopsis* as the backbone on which to map participating gene copies and their variants [18], making different cultivars and species more easily comparable. As a result, we established similar patterns of variation in the Jersey kale genome as previously have been found in the ‘Kashirka’ cultivar. However, with this approach, the importance of additional genes outside the homologous pathway is never discovered. In contrast, our approach also uncovered the role of Bo6G120900 (TWIN SISTER OF FT, TSF). As such, the technique we present here is at least complementary, and especially useful in exploring less well-characterized pathways and discovering participating genes.

## Conclusions

We performed a bulk segregant analysis (BSA) across four cohorts of a cross between the Jersey kale and the *B. oleracea* model TO1000DH3 phenotyped on flowering time. The data we collected consisted of high throughput sequencing reads, which we analyzed using standard tools for identifying QTLs in pairwise BSA comparisons. This resulted in numerous regions throughout the genome, though concentrated at loci known from previous research in flowering time regulation and consistent across pairwise comparisons. To reduce the set of candidate loci to more manageable dimensions, we developed an approach that limits the result set by focusing on genes annotated with terms contained within relevant subgraphs of the Gene Ontology. This reduced the resulting gene set from tens of thousands to dozens of candidate genes. A further enrichment test led to the pathway for circadian rhythm in plants.

The genes that enriched this pathway are attested from previous research as being involved in regulating flowering time, and some of these genes were also identified as having functionally significant variation compared to *Arabidopsis*. As such, we validated and confirmed our ontology-mediated results through a more targeted, homology-based approach. However, the ontology-mediated approach produced additional genes of putative importance, showing that the approach aids in exploration and discovery. We view our method as potentially applicable to the study of other complex traits and therefore make our workflows available as open-source code and a reusable Docker container.

## Data availability

- Sequencing data produced in this project are deposited at SRA under project number PRJNA564368.
- The supplementary tables referred to in the manuscript body are contained in a (CC4-Attribution licensed) archive that additionally holds an extended methods section with links to all other inputs and outputs (e.g. raw, intermediate, and result data) of this project. This archive is identified by 10.5281/zenodo.3402201
- The analytical approach we describe in this manuscript is made available as open source code (MIT license) identified by 10.5281/zenodo.5211374 and also as a Docker container that additionally has all required 3^rd^ party tools for performing the analyses described in this manuscript. The container is identified on the Docker hub by naturalis/brassica-snps
- The SnpEff analyses and their results have been recorded separately in an archive identified by 10.5281/zenodo.5211461

## Acknowledgements

The authors are grateful to Sarah-Veronica Schießl-Weidenweber for the insightful exchanges we had about her previous work on flowering time in *B. oleracea* and for the background data she provided on genomic coordinates of salient genes and their features. Lab support for DNA sequencing was provided by Elza Duijm of Naturalis. The SnpEff analysis was performed by Esther Kockelmans, Rik Frijmann, Nino Vrolijk and Daphne van Ginneken of the University of Applied Sciences Leiden. This work was supported by Naturalis Biodiversity Center [grant number 077 LENS].

